# Optogenetic quantification of source sink relationship in intact hearts to explain cardiac arrhythmia initiation and protection

**DOI:** 10.1101/2024.08.14.604123

**Authors:** Judith S. Langen, Patrick M. Boyle, Daniela Malan, Philipp Sasse

## Abstract

Increased cardiac excitability and reduced electrical coupling promote cardiac arrhythmia and can be quantified by input resistance (R_m_), pacing threshold (I_thr_), and cardiac length constant (λ). However, measurement of these parameters in the heart was not feasible, because the required homogenous current injection cannot be performed with electrical stimulation. Here, we overcame this problem by optogenetic current injection into all illuminated cardiomyocytes of mouse hearts in different action potential phases. Precisely triggered and patterned illumination enabled measuring R_m_ and λ, which both were smallest at diastole and larger during plateau and repolarization. Pharmacological and depolarization-induced reduction of inward rectifying K^+^ currents (I_K1_), gap junction block and cardiac infarction reduced I_thr_ showing the importance of high I_K1_ density and intact cardiomyocyte coupling for preventing arrhythmia initiation. Simulations in a calibrated cardiomyocyte model were used to classify pro- and anti-arrhythmic mechanisms based on their effects on R_m_ and I_thr_. Finally, combining experiments with simulations allowed for quantification of I_K1_ inward rectification in the intact heart, identifying strong rectification as a new pro-arrhythmic concept.

## Introduction

Due to electrical coupling of cardiomyocytes through gap junctions, the heart is a functional electrical syncytium, which is important for rapid conduction of electrical excitation but also for prevention of pathological activity. In cardiac tissue, excitatory impulses act as sources of depolarizing current for adjacent repolarized tissue (sink) and the source current density must be sufficient to depolarize the sink to its activation threshold or propagation will fail^1^. Consequently, in well-coupled healthy myocardium a pathological afterdepolarization in an individual cardiomyocyte (source) results in electrotonic current flow to the neighboring repolarized cardiomyocytes (sink), failing to induce a potentially dangerous premature ventricular contraction (PVC)^2^. Thus, loss of coupling plays a critical role in arrhythmogenesis, not only by slowing conduction, but also by reducing the protective sink and, accordingly, decreasing the source current density required to trigger a PVC^1^. Additionally, repolarizing outward currents influence cardiac electrical stability by amplifying sink effects and counteracting potential afterdepolarizations. At rest, the major contributor to these outward currents is the inwardly rectifying K^+^ current (I_K1_), which stabilizes resting membrane potential (RMP) and shapes final repolarization^3^. Its outward K^+^ current is regulated by voltage- dependent block of the pore by intracellular positively charged polyamines, resulting in a reduced conductance at depolarized membrane potentials^3^, which is important for fast action potential (AP) upstroke and conduction velocity.

Ventricular arrhythmia due to afterdepolarization-induced PVCs is the leading cause of sudden cardiac death^4^. Quantification of cardiac excitability and cell-to-cell coupling in the intact heart is crucial for investigating mechanisms of arrhythmia initiation and maintenance. Three separate but linked parameters are important to consider: 1) The input resistance (R_m_) of a cardiomyocyte, which describes the membrane potential change in response to a subthreshold current injection. Low diastolic R_m_ is attributed to the high conductance of I_K1_. Thus, if a pathological depolarizing current arises, the resulting (after-) depolarization, given mathematically by Ohm’s law as the product of R_m_ and current amplitude, will be small. However, if I_K1_ is pathologically reduced, as in heart failure^5,6^, R_m_ increases and the voltage change caused by the same current will be larger, potentially causing a PVC^7^. 2) The pacing threshold as a measure of the current density sufficient to trigger an AP in a single cell (I_thr_). 3) The length constant (λ), which describes the spread of subthreshold depolarization in space via electrotonic current flow to neighboring cardiomyocytes, quantifying electrical coupling.

Until recently, it was not possible to precisely measure R_m,_ I_thr_, and λ in the intact heart because homogenous depolarization of cardiomyocytes is required, but was neither feasible by single cell current injection in the syncytium of electrically coupled cardiomyocytes nor by extracellular electrical stimulation. Thus, previous approaches were limited to isolated cells and small tissue preparations or used non-homogeneous extracellular field stimulation, which induces areas of hyperpolarization^8^. To overcome these limitations, we have developed optogenetic current clamp to directly quantify R_m_, I_thr_, and λ in the intact heart with high temporal resolution during the cardiac cycle. This is enabled by precisely timed, confined illumination of hearts expressing the light-gated cation channel channelrhodopsin-2 (ChR2), which leads to the required current injection in all illuminated cardiomyocytes^9^.

Because many pathological conditions, including heart failure and myocardial infarction, involve increased occurrence of cardiac arrhythmias^5,10^, we propose that quantification of susceptibility to afterdepolarizations and PVCs as well as electric coupling in intact hearts is important to improve our understanding of arrhythmia mechanisms and to develop new therapeutic concepts.

## Results

### Optogenetic determination of R_m_ in intact hearts

We developed an optogenetic current clamp method to measure R_m_ in the intact heart and in different AP phases. Precisely timed current injection in all cardiomyocytes of the left ventricular free wall was performed by epicardial illumination of Langendorff-perfused mouse hearts expressing the light-gated ion channel channelrhodopsin-2^9^. High intensity light pulses paced hearts with a fixed cycle length of 275 ms followed by low intensity subthreshold light pulses with defined delays (**Fig. 1a,b**). Membrane potential was measured by sharp microelectrode recording and averaged over multiple cycles to reduce noise (**Fig. 1c**). Subthreshold depolarization (ΔE) was calculated as difference between the averaged APs with and without subthreshold illumination (**Fig. 1d,g**). The injected ChR2 current (I_ChR2_) was computed by a ChR2 gating model^11^ (**Fig. 1e,h**), which was calibrated to match measurements from patch clamp experiments of ventricular cardiomyocytes expressing ChR2 (**Fig. 1f**, n = 7). R_m_ was calculated according to Ohm’s law as R_m_ = ΔE/I_ChR2_ for each delay (**Fig. 1i,j**), and was significantly smaller during diastole (63.3±13.1 MΩ = 100%) compared to the plateau (137.4±10.3%), early (APD70, 227.9±11.2%), and late (APD90, 133.4±3.9%) repolarization. Furthermore, R_m_ at APD70 was significantly higher than R_m_ during other AP phases suggesting that the heart is particularly vulnerable to afterdepolarizations triggering PVCs in this early repolarization phase.

**Figure 1:**
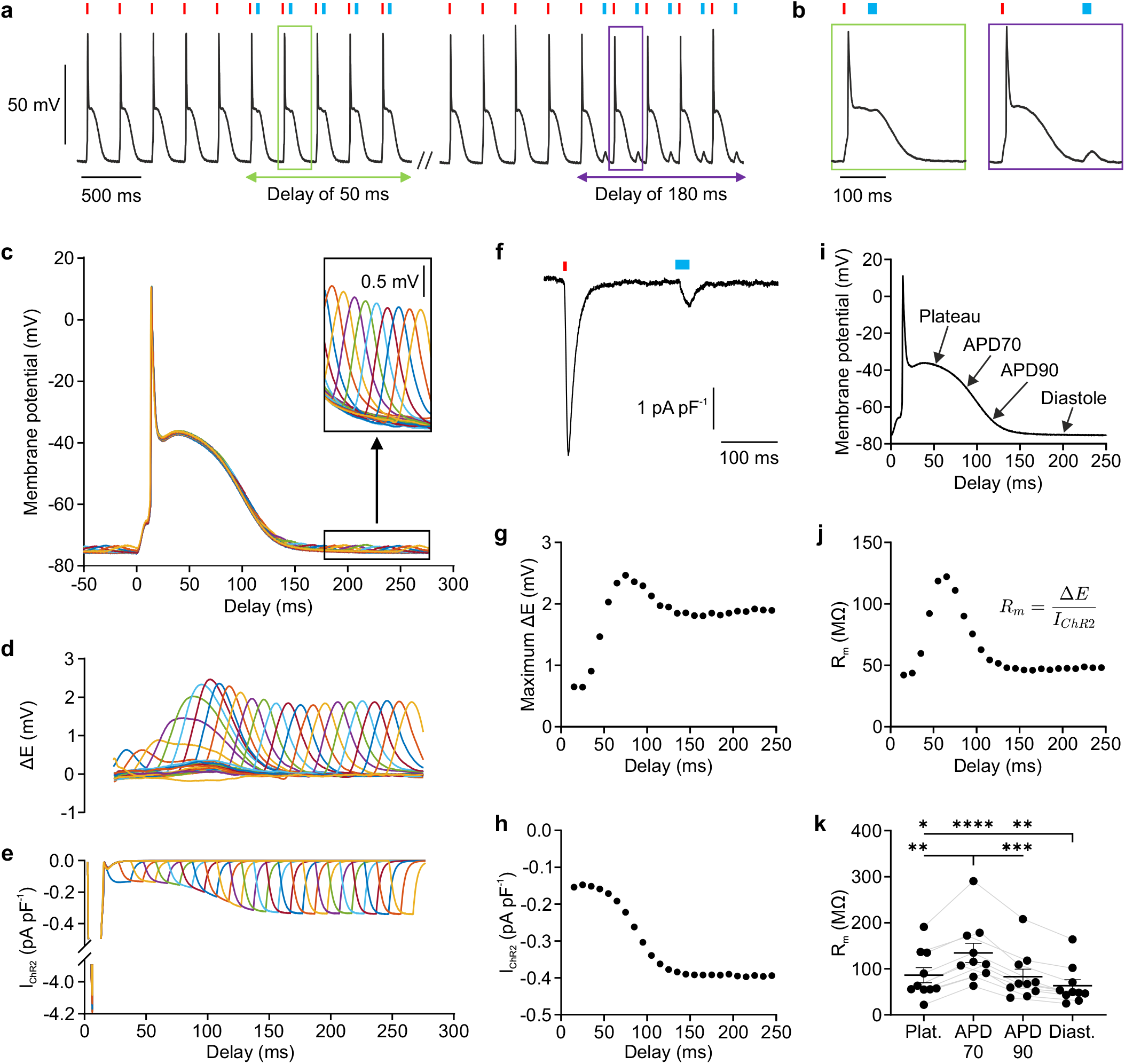
Optogenetic determination of input resistance (R_m_) in intact hearts. **a,b**, Representative AP trace with pacing (red, ∼350 µW mm^-2^, 5-10 ms) and subthreshold (blue, 15-40 µW mm^-2^, 20 ms) light stimulation (465 nm). **c**, Averaged APs with second subthreshold light pulses of delays between 10 and 240 ms. **d,e**, Membrane potential change (**d**, ΔE) calculated as difference between averaged APs with and without subthreshold light pulse and corresponding ChR2 current (**e**, I_ChR2_) computed by a ChR2 gating model for the different delays. **f**, ChR2 current of a ventricular cardiomyocyte evoked by high (red, 315.5 µW mm^-2^) and subthreshold (blue, 13.6 µW mm^-2^) illumination analogue to the stimulation in (a-e). **g**-**j**, Representative averaged AP (**i**) and corresponding values of R_m_ (**j**) calculated as ratio of maximum membrane potential change (**g**) and ChR2 current (**h**) (a-j from one representative heart or cardiomyocyte). (**k**) R_m_ at AP plateau, 70% and 90% of repolarization (APD70 and APD90) and diastole (repeated measures one-way ANOVA, Tukey’s multiple comparisons post-test, N = 10, p < 0.0001). Mean ± SEM. *p < 0.05, **p < 0.01, ***p < 0.001, ****p < 0.0001.

### Quantification of I_K1_ contribution to R_m_ and I_thr_

To determine the contribution of I_K1_ to R_m_ in different AP phases, the I_K1_ blocker BaCl_2_ was applied at low concentration, resulting in a small AP prolongation (**Fig. 2a**) and an increase in diastolic R_m_ of 23.6±4.6% (**Fig. 2b,d**), but without affecting R_m_ during early repolarization (APD70, **Fig. 2c**). Interestingly, this concentration of BaCl_2_ did not depolarize cells (**Fig. 2e**) or change the maximum AP upstroke velocity (**Fig. 2f**). To show that I_K1_ reduction can underlie the vulnerability of the heart to afterdepolarizations, we analyzed the amplitude of spontaneous diastolic delayed after-depolarizations (DADs). DADs were promoted by fast pacing to induce Ca^2+^ overload in the intracellular stores, which alone did not produce DADs (**Fig. 2g,h**, black). However, in combination with caffeine, which sensitizes ryanodine receptors to release Ca^2+^, spontaneous DADs of an amplitude of 1.23±0.25 mV (**Fig. 2g,h**, red) could be observed. Administration of BaCl_2_ increased DAD amplitude to 1.87±0.27 mV (**Fig. 2g,h**, blue). Calculation of the currents underlying these DADs using Ohm’s law (I_DAD_ = E_DAD_ / R_m_) showed almost identical values (caffeine: 32.9 pA, caffeine + BaCl_2_: 33.2 pA). Thus, the change in DAD amplitude could be completely explained by the increased R_m_.

**Figure 2:**
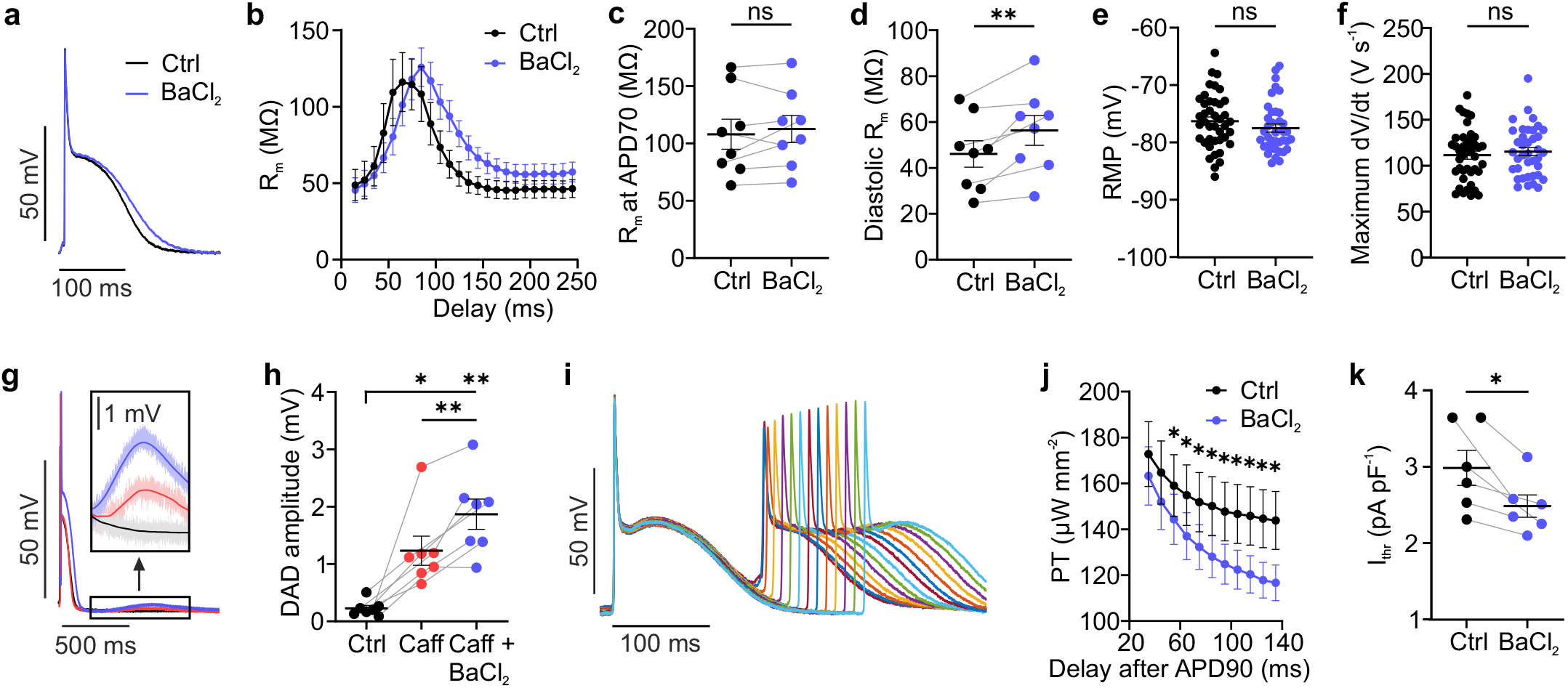
Effect of I_K1_ reduction on cardiac excitability. **a,b**, Representative, averaged APs (**a**) and R_m_ for different delays (**b**) of the subthreshold light pulse with and without BaCl_2_ (10 µM). **c**-**f**, Quantification of the effect of I_K1_ block on R_m_ during early repolarization (**c**, at APD70, two-tailed, paired t-test, N = 8, p = 0.39) and diastole (**d**, two-tailed, paired t-test, N = 8, p = 0.0016), RMP (**e**, two-tailed, unpaired t-test, N = 4, n = 38-41, p = 0.25) and maximum upstroke velocity (**f**, two-tailed, unpaired t-test, N = 4, n = 38-41, p = 0.53). **g**, DADs resulting from fast pacing (cycle length 70-140 ms) without (black), with caffeine (1 mM, red) and with BaCl_2_ additionally to caffeine (blue). **h**, Statistical analysis of DAD amplitude (repeated measures one-way ANOVA, Tukey’s multiple comparisons post-test, N = 7, p = 0.0003). **i**, Representative AP traces of a S1-S2 protocol (last S1showed) followed by a premature stimulus (S2) of variable delay (120-260 ms). **j,k**, Thresholds for optogenetic pacing (PT) for different delays of S2 (10 ms binning) after APD90 (**j**) and statistics of diastolic I_thr_ determined 135 ms after APD90 (**k**, two-tailed, paired t-test, N = 6, p = 0.02) with and without BaCl_2_. Mean ± SEM. *p < 0.05, **p < 0.01, ns, not significant.

To investigate if the increased R_m_ also corresponds with an increased susceptibility to PVCs, we determined the threshold for optogenetic pacing in different AP phases (**Fig. 2i**) and calculated diastolic I_thr_, which was significantly reduced by 15.77±3.42% in the presence of BaCl_2_ (**Fig. 2k**). Beyond I_K1_ reduction, depolarization of the RMP has been discussed as another critical factor promoting cardiac arrhythmia in ischemic and failing hearts^5,6^. To investigate the effect on R_m_ and I_thr_, sustained low-intensity illumination was applied during diastole (**Fig. 3a**, light blue bar), which increased RMP up to ∼10 mV. Additionally, short pacing and subthreshold light pulses were applied to determine I_thr_ and R_m_, respectively (**Fig. 3a**, red and dark blue bar). RMP depolarization significantly increased diastolic R_m_ (**Fig. 3b,c**) and decreased I_thr_ (**Fig. 3d,e**).

**Figure 3:**
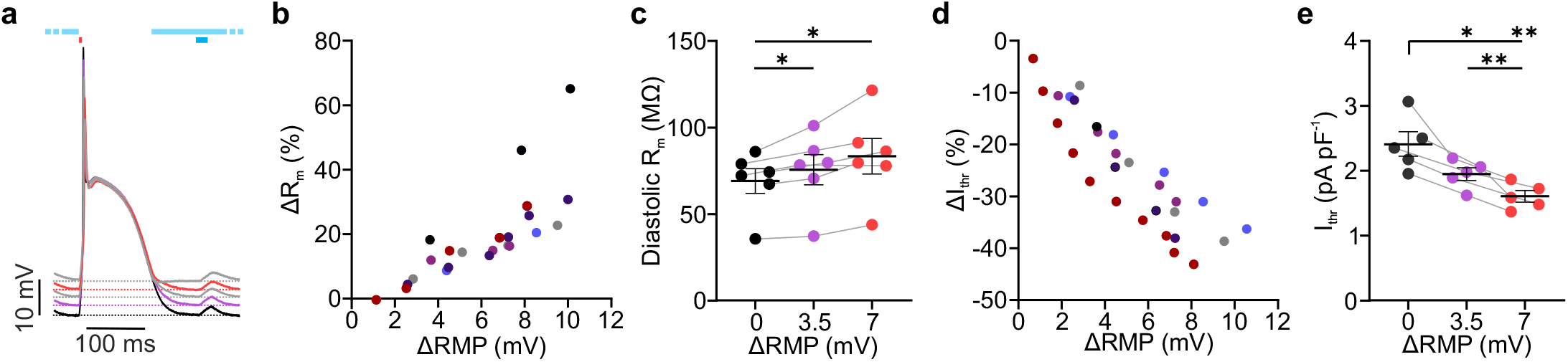
Effect of RMP depolarization on cardiac excitability. **a**, Representative, averaged, paced (red bar) AP traces without (black) and with low-intensity illumination during the diastole (9.4-33.6 µW mm^-2^, light blue bar) and subthreshold pulses to determine R_m_ (dark blue bar). **b**-**e**, Relationship between depolarization of RMP (ΔRMP) and change of R_m_ (**b**, ΔR_m_) or I_thr_ (**d**, ΔI_thr_) with different colors indicating individual hearts. Statistical analysis of diastolic R_m_ (**c**, N = 6, p = 0.021) and I_thr_ (**e**, N = 5, p = 0.0015) for 3.5 and 7 mV RMP depolarization (repeated measures one-way ANOVA, Tukey’s multiple comparisons post-test). Mean ± SEM. *p < 0.05, **p < 0.01.

### Modelling of ionic currents underlying cardiac excitability

To investigate the relative contributions of individual currents to cardiac excitability and RMP in the diastole, we conducted computer simulations allowing for modification of ion current densities and I_K1_ inward rectification. The simulations were performed using a model of a mouse ventricular cardiomyocyte^12^ incorporating the ChR2 gating model^11^. To quantify RMP, R_m_, and I_thr_ in the model, an analysis framework was developed in openCARP to mimic experimental conditions. We found that decreasing I_K1_ or Na^+^/K^+^-ATPase current (I_NaK_) as well as increasing Na^+^ background current (I_Nab_) led to depolarization of the RMP (**Fig. 4a**), increase in R_m_ (**Fig. 4b**) and decrease in I_thr_ (**Fig. 4c**) and vice versa. Interestingly, an increase in I_K1_ inward rectification (ir) led to similar effects on R_m_ and I_thr_ without affecting the RMP (**Fig. 4a-c**).

**Figure 4:**
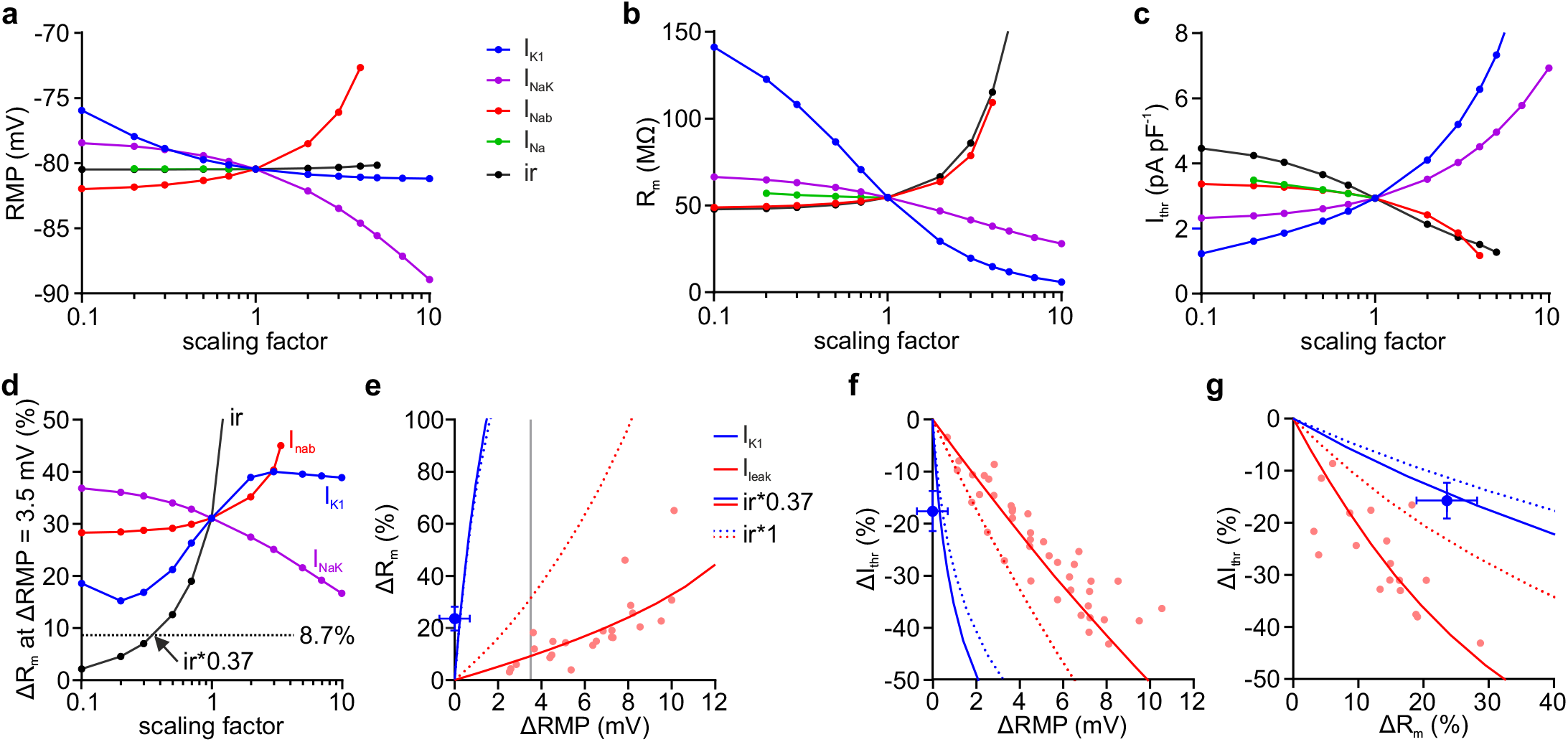
Contribution of diastolic ion currents to excitability in a model of a mouse ventricular cardiomyocyte. **a**-**c**, Effect of scaling I_K1_, Na^+^/K^+^-ATPase current (I_NaK_), sodium background current (I_Nab_), voltage-dependent sodium current (I_Na_), and inward rectification of I_K1_ (ir) on RMP (**a**), diastolic R_m_, (**b**) and I_thr_ (**c**). **d**, Relative change in R_m_ in response to RMP depolarization of 3.5 mV by adding a cation background current (I_leak_) mimicking I_ChR2_ for different scaling factors of diastolic currents. Dotted, black line indicates the experimentally measured increase in R_m_ of 8.7%. **e**-**g**, Relationships between relative changes in RMP, R_m_, and I_thr_ predicted by the model with ir*1 (dotted lines) and ir*0.37 (solid lines) for reduction of I_K1_ (blue) and increase in I_leak_ (red). Grey line indicates effect of 3.5 mV RMP depolarization on R_m_ (**e**). Experimental data from Fig. 2 (blue, mean ± SEM) and 3 (red) are shown as dots.

To compare experimental data with simulation predictions, we modeled the effect of RMP depolarization by adding a cation background current (I_leak_) mimicking I_ChR2_. Experimentally, a RMP depolarization of 3.5 mV led to an increase in R_m_ of 8.7% (**Fig. 3b,c**) and in simulations, the same RMP depolarization resulted in an R_m_ increase of ∼30% (**Fig. 4e**, intersection of dashed red and grey line). To reduce this discrepancy, we tested scaling of the currents involved during diastole (**Fig. 4d**). Importantly, only a reduction of the I_K1_ inward rectification (ir) to 37%, but not up- or downscaling of current densities, was able to explain the experimental depolarization-induced change in R_m_ (**Fig. 4d**, dotted, black line). Moreover, the calibrated (**Fig. 4e-g**, ir*0.37, solid lines), but not the original model (**Fig. 4e-g**, ir*1, dotted lines), perfectly predicted the relationships between changes of RMP, R_m_, and I_thr_ of the experimental data (dots) from optogenetic RMP depolarization (red) and I_K1_ reduction by BaCl_2_ (blue). Furthermore, the interdependency of these three excitability parameters can be used to discriminate between increased excitability due to I_K1_ block, which reduced I_thr_ and increased R_m_ without affecting RMP (**Fig. 4e-f, blue**), and increased excitability due to increased leak current, which additionally depolarizes RMP (**Fig. 4e-f**, red) and has a stronger effect on I_thr_ than on R_m_ (**Fig. 4g**). Patients with pro-arrhythmic increase in cardiac excitability can benefit from sodium channel blockers like Lidocaine. As expected, Lidocaine led to a reduction of the AP upstroke velocity and amplitude (**Fig. 5a**, insert) and significantly reduced I_thr_ (**Fig. 5b**) without affecting R_m_ (**Fig. 5c**). This was confirmed in simulations, which showed a similar increase in I_thr_ (**Fig. 5d,e**, green) and only a small increase in R_m_ (**Fig. 5d,f**, green). Additionally, we simulated effects of the anti-arrhythmic drugs Amiodarone and Dronedarone, which block sodium currents but also affect repolarizing K^+^ currents, Ca^2+^ currents, and the Na^+^/K^+^-ATPase, using the degree of block reported in Loewe et al^13^. These simulations showed that Dronedarone behaved similar to Lidocaine with minor effects on R_m_, whereas Amiodarone increased R_m_ (**Fig. 5f**). This suggests a relevant block of stabilizing K^+^ conductances, which explains the smaller effect on I_thr_ compared to the other two blockers (**Fig. 5e**).

**Figure 5:**
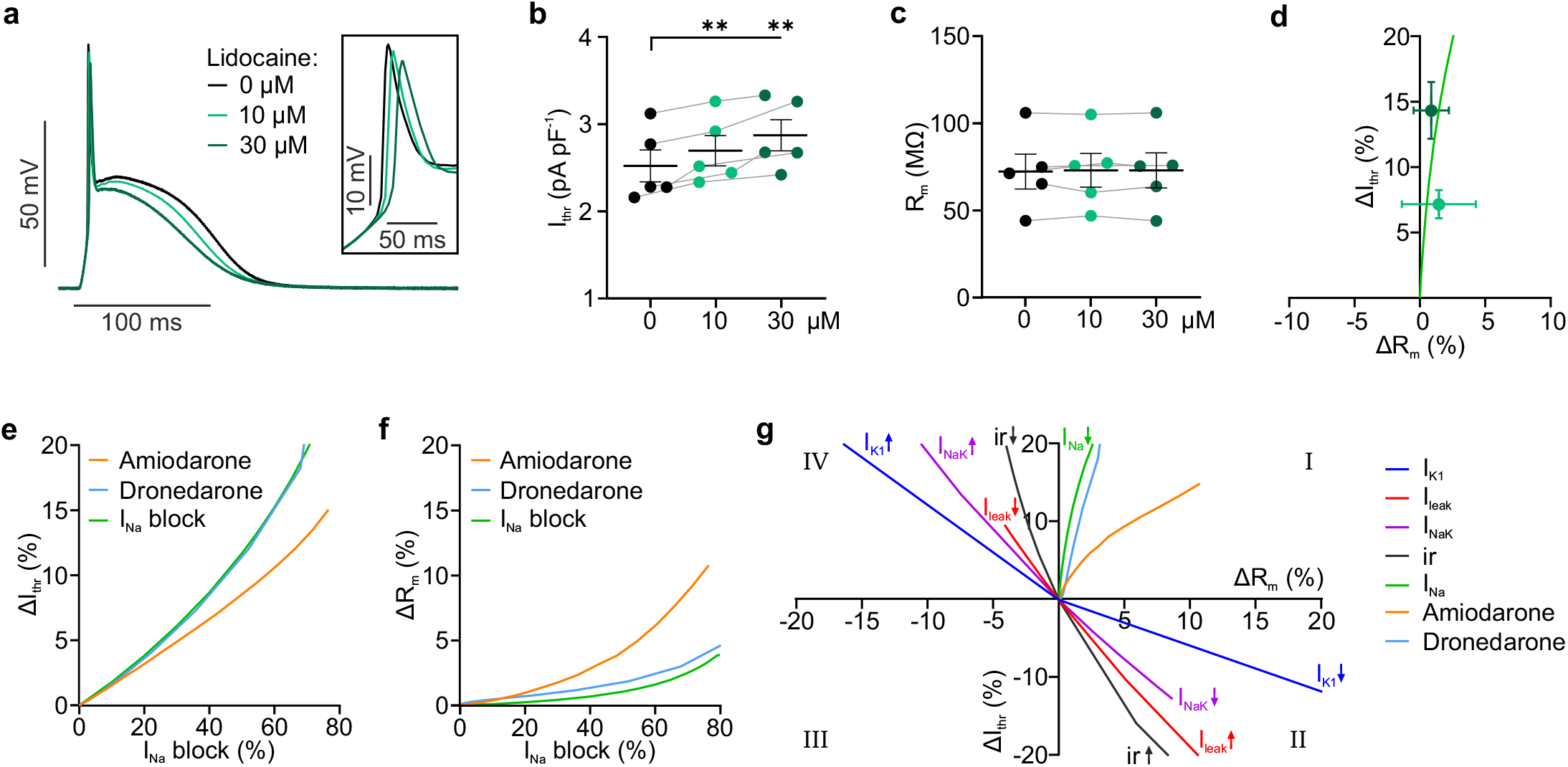
Effect of Na^+^ and multi-channel blockers on cardiac excitability. **a-c**, Representative, averaged APs (**a**), I_thr_ (**b**, N = 5, p = 0.0022) and diastolic R_m_ (**c**, N = 5, p = 0.73) without and with the sodium channel blocker Lidocaine (10, 30 µM, repeated measures one-way ANOVA, Tukey’s multiple comparisons post-test). **d**, Relationship between change of R_m_ (ΔR_m_) and I_thr_ (ΔI_thr_) resulting from sodium current reduction in simulations (solid line) and experiments (dots). **e**-**f**, Simulated relationship between degree of I_Na_ block and increase in pacing threshold (**e**) and R_m_ (**f**) for increasing concentrations of the multi-channel blockers Amiodarone and Dronedarone and isolated I_Na_ block. **g**, Summary of the relationship between ΔR_m_ and ΔI_thr_ for isolated modifications of indicated ion channels and the multi-channel blockers from simulations in the calibrated cardiomyocyte model. Mean ± SEM. **p < 0.01.

Our experiments suggest that the interdependency of R_m_ and I_thr_ is highly specific for different blockers (**Fig. 4g, 5d**). To classify effects of drugs or pathological conditions affecting cardiac excitability, we simulated their effects on the relationship between change in R_m_ and I_thr_ (**Fig. 5g**, quadrant I - IV). Quadrant I describes the effect of multi-channel blockers, which reduce excitability by blocking Na^+^ channels, thus increasing I_thr_; this effect may be counteracted by blocking K^+^ currents, leading to increased R_m_ (e.g. Amiodarone). Quadrant II shows pathological situations with reduced I_K1_ or Na^+^/K^+^-ATPase as well as increased leak currents or I_K1_ inward rectification leading to an increase in R_m_ and decrease in I_thr_. These effects could be reversed by the antiarrhythmic conditions shown in quadrant IV which will protect from PVCs not only by increasing I_thr_ but also by decreasing R_m_.

### Optogenetic determination of λ in the intact heart

To quantify electrical coupling between cardiomyocytes in the intact heart, we developed an optogenetic method to measure λ by localized subthreshold stimulation. First, a mathematical description of the magnitude of depolarization depending on the distribution of light and the electrical coupling was derived. Without electrical coupling, the regional distribution of depolarization in response to optogenetic stimulation equals the distribution of light (**Fig. 6a**, blue function *f*). However, electrical coupling between the cells leads to a sigmoidal distribution of depolarization along the boundary of the illumination area (**Fig. 6a**, red function *f* * *g*), which can be explained by electrotonic current flow between the adjacent cells. The resulting magnitude of depolarization in space can be described by convolution of the light function *f* with a weight function *g*, which describes the exponential spread of a subthreshold stimulus in one dimension and depends on λ (**Fig. 6a**).

**Figure 6:**
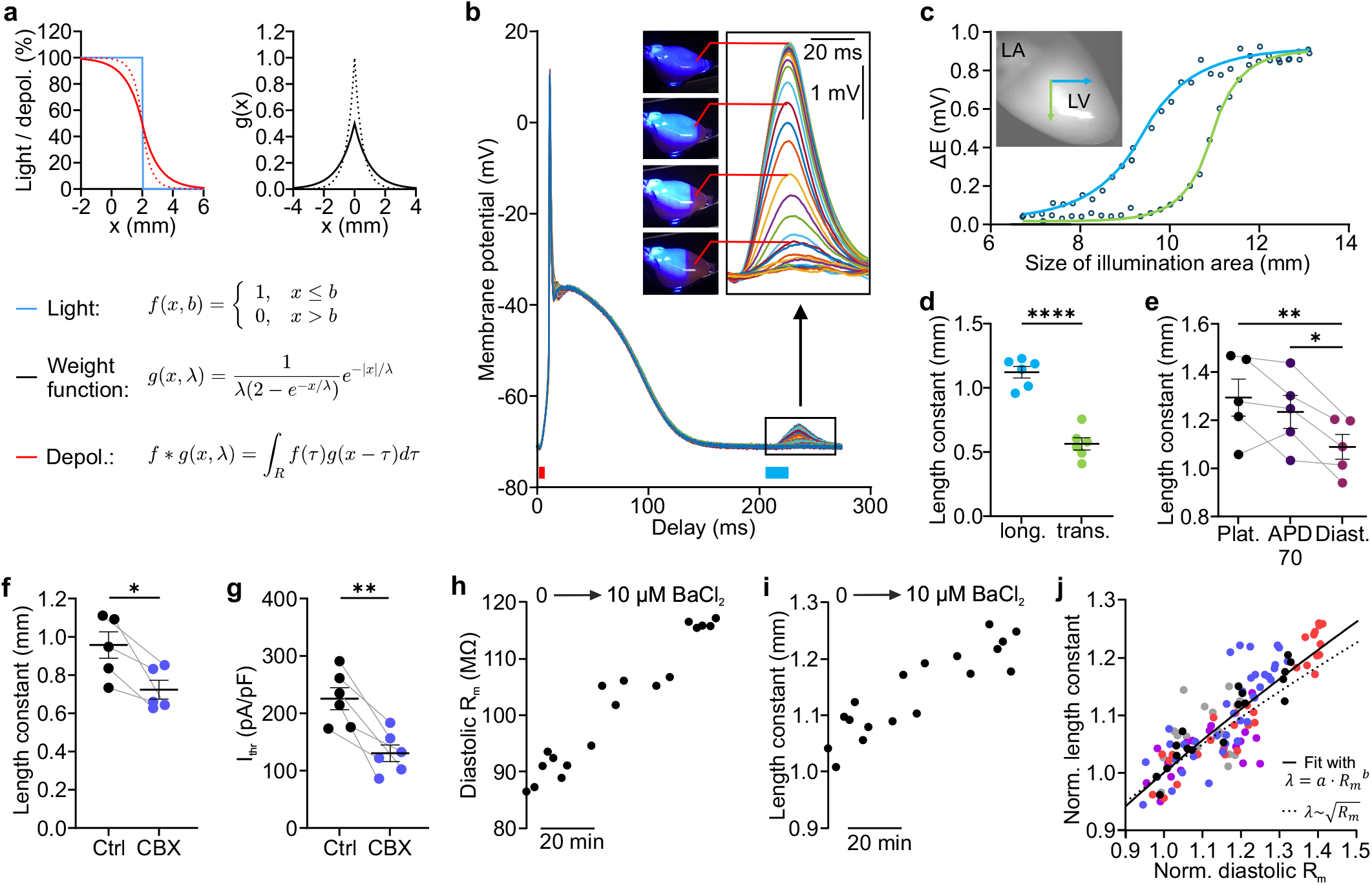
Optogenetic determination of cardiac length constant (λ) in the intact heart. **a**, Concept for determination of λ. Theoretical example of depolarization in 1-dimensional space (left) resulting from illumination restricted in space (blue) with high (solid red, 1 mm) and low (dotted red, 0.5 mm) λ. The magnitude of depolarization is calculated by convolution (*f* * *g*) of the distribution of light (*f*) with a weight function (*g*, right) which depends on λ (solid: 1 mm, dotted: 0.5 mm). **b**, Averaged APs with different illumination sizes of the subthreshold light pulses (blue bar) applied after a global pacing pulse (red bar). **c,d**, Relationship between subthreshold membrane potential change (ΔE) and size of illumination area fitted with the convolution function *f* * *g* (**c**) and diastolic λ obtained from fitting λ as parameter of *g* upon decreasing the illumination area in longitudinal (blue) and transversal (green) direction of fiber orientation (**d**, two-tailed, unpaired t-test, N = 6, p < 0.0001). **e**, Longitudinal λ at plateau (Plat.), 70% of repolarization (APD70), and diastole (Diast., repeated measures one-way ANOVA, Tukey’s multiple comparisons post-test, N = 5, p = 0.008). **f,g**, Diastolic, longitudinal λ (**f**, N = 5, p = 0.032) and I_thr_ (**g**, N = 6, p = 0.0028) without and with the gap junction blocker Carbenoxolone (CBX, 10 µM, two-tailed, paired t-test). **h**-**j**, Change in diastolic R_m_ (**h**) and λ (**i**) during successive increase of BaCl_2_ concentration up to 10 µM in one representative heart and normalized changes in R_m_ and λ (**j**) fitted with a power function (solid black, a = 1.002, 99% confidence interval: [0.986,1.018], b = 0.572, 99% confidence interval: [0.487, 0.656]). Square root relationship shown with dotted line. Colors indicate different hearts. Mean ± SEM. *p < 0.05, **p < 0.01, ****p < 0.0001.

Experimentally, λ was determined by successively decreasing the illumination area, which led to an increased distance to the recording site (**Fig. 6b**, insert, recording site in red), and therefore, to a reduction of the subthreshold depolarization (**Fig. 6b,c**). Because λ depends on the fiber orientation^14^, we measured λ in longitudinal and transverse fiber directions (**Fig. 6c**, blue and green arrows, respectively). Fitting the relationship between size of illumination area and magnitude of subthreshold depolarization with the convolution function *f* * *g* (**Fig. 6c**) could be used to reliably determine λ, which was significantly larger in longitudinal (**Fig. 6d**, blue, 1.12±0.05 mm) than in transversal direction of fiber orientation (**Fig. 6d**, green, 0.56±0.05 mm). Additionally, we measured the longitudinal λ at different timepoints during the cardiac cycle and found that the diastolic λ (**Fig. 6e**, 1.09±0.05 mm) was significantly smaller compared to λ at early repolarization (**Fig. 6e**, 1.23±0.07 mm) or plateau (**Fig. 6e**, 1.29±0.08 mm). To validate this method, we applied the gap junction blocker Carbenoxolone which significantly reduced diastolic λ by 22.3±5.7% (**Fig. 6f**). Carbenoxolone also decreased I_thr_ by 41.3±6.2% (**Fig. 6g**), suggesting a critical role of uncoupling in arrhythmia initiation.

Cable theory predicts a square root relationship between λ and 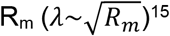, but this has not been experimentally proven in intact hearts. Thus, we applied increasing concentrations of the I_K1_ blocker BaCl_2_ and measured the increase in diastolic R_m_ and λ at identical timepoints (**Fig. 6h,i**). Their relationship was quantified by fitting with a power function (**Fig. 6j**, solid black line) and was consistent with the square root relation (dotted black line).

### Effect of electrical uncoupling on cardiac excitability

Myocardial infarcts (MI) increase the risk of PVCs triggering potential lethal ventricular arrhythmia. To quantify the mechanisms involved, we determined the PVC vulnerability after acute cardiac infarction in the remote, healthy myocardium (**Fig. 7a**, blue), the infarcted area with loss of cardiomyocytes (**Fig. 7a**, purple), and the border zone in between (**Fig. 7a**, red). PVC vulnerability was calculated as the reciprocal of the threshold for localized optogenetic pacing determined by successively moving the illumination pattern from the remote to the infarction (**Fig. 7a**). After infarction, PVC vulnerability was significantly increased near the border of infarction (**Fig. 7b**, pattern -3 to -1) and decreased in the infarcted area (pattern 1, 2, and MI). Furthermore, after infarction the PVC vulnerability was significantly higher in the border zone compared to the remote myocardium (**Fig. 7d,e**, +47.6±8.6%), whereas there was no significant difference between the two locations before infarction (**Fig. 7c**). Because increased PVC vulnerability could be due to loss of electrical sink, but also due to RMP depolarization (**Fig. 3**), we measured RMP at different distances from the infarcted area. Near the border of the infarct, RMP was depolarized up to 3.5 mV (although not significant) compared to RMP before infarction (**Fig. 7f**), which would increase PVC vulnerability by 18.4±2.6% (**Fig. 3d,e**). This indicates that RMP depolarization accounts for <50% of the increased PVC vulnerability of 47.6±8.6% in the border zone and >50% must be because of a reduced electrical sink due to loss of cardiomyocytes in the infarcted area.

**Figure 7:**
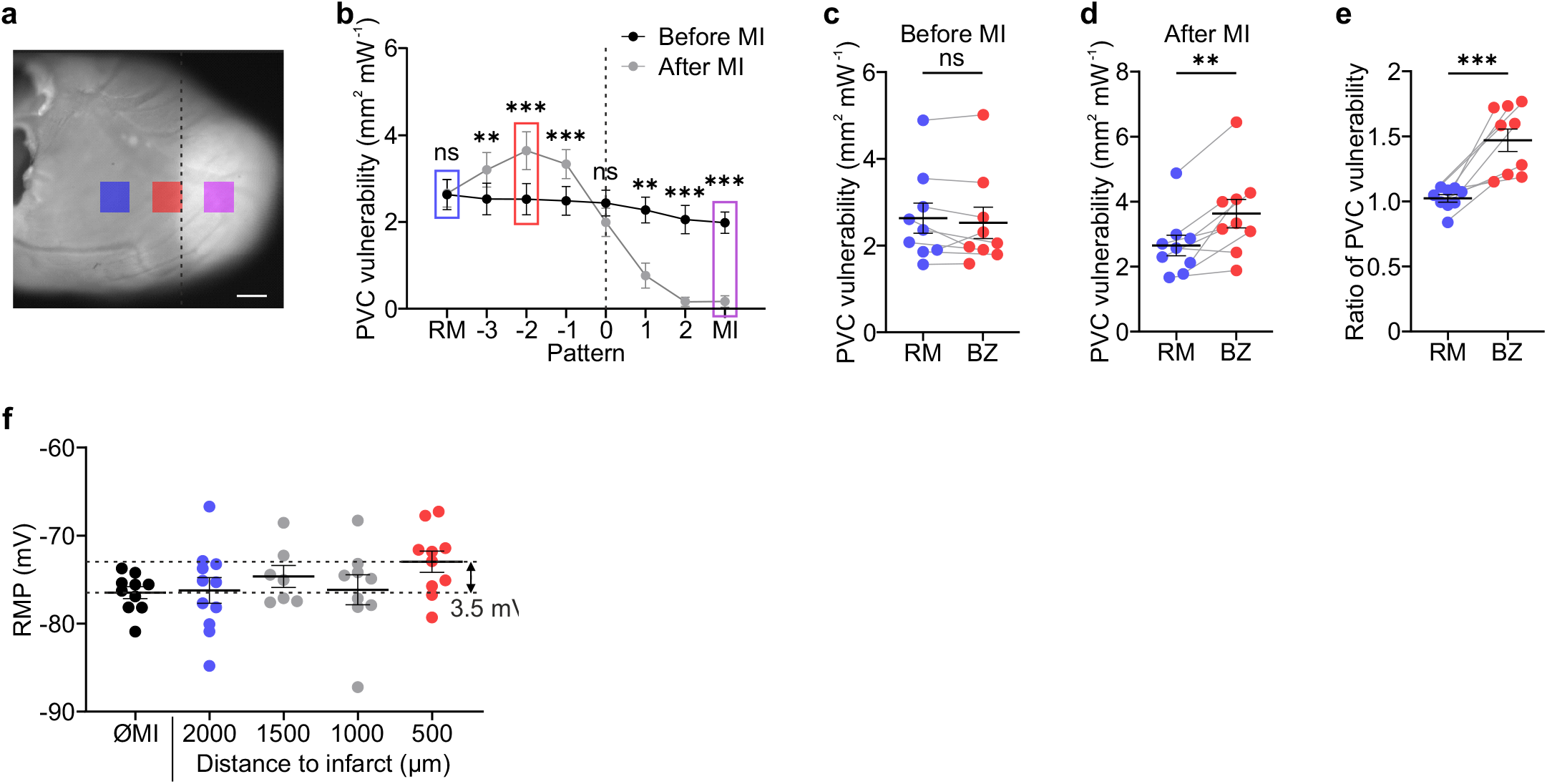
Increased cardiac excitability after acute myocardial infarction. **a,b**, Image of the left ventricle (**a**) after cryoinfarction (white) at the heart apex (scale bar: 1 mm) and PVC vulnerability defined as reciprocal of pacing threshold before and after infarction for different regions (**b**). The border of infarction is indicated with dashed line and pattern of remote myocardium (RM), border zone (BZ, pattern -2), and infarction (MI) are highlighted in blue, red, and purple, respectively (**a,b**). **c**-**e**, Statistical analysis of PVC vulnerability before (**c**, two-tailed, paired t-test, N = 9, p = 0.3) and after infarction (**d**, two-tailed, paired t-test, N = 9, p = 0.0025) and the ratio of PVC vulnerability after and before infarction (**e**, two-tailed, paired t-test, N = 9, p = 0.0005). **f**, RMP before (∅MI) and after cryoinfarction recorded at different distances from the border of infarction (Ordinary one-way ANOVA, N = 2-3, n = 9-11, p = 0.27). Mean ± SEM. **p < 0.01, ***p < 0.001, ns, not significant.

## Discussion

Reduced electrical coupling and increased excitability are important factors in myocardial infarction, heart failure, arrhythmia, and sudden cardiac death. Previous studies focused on disease-specific changes in currents by patch clamp and expression analysis. However, the changes in any individual channel or current can be small and their interaction complex, making the consequences for arrhythmia difficult to predict. We suggest that quantification of functional parameters such as R_m_, I_thr_, and λ in the intact heart is essential for detailed understanding of electrophysiological changes that promote arrhythmias.

Our light-induced current clamp method enabled to determine these functional arrhythmia parameters in the intact heart. The diastolic R_m_ in mouse hearts (∼60 MΩ, **Fig. 1k**) was in the range of those measured by current injection in isolated rat (40-60 MΩ)^16,17^ and guinea pig (∼10 MΩ)^18^ ventricular cardiomyocytes. However, R_m_ in canine papillary muscle (20-225 kΩ)^19^ or tissue islands of this muscle (0.25-1.5 MΩ)^19^ was much smaller, presumably because part of the injected current was lost due to electrotonic current flow to neighboring cardiomyocytes, leading to R_m_ underestimation. In contrast, optogenetic stimulation allows for light-induced current injection in all cardiomyocytes expressing ChR2.

The use of patterned, optogenetic current clamp and mathematical convolution allowed to precisely determine λ in longitudinal (∼1.1 mm) and transversal (∼0.6 mm) direction of fiber orientation (**Fig. 6d**), which is in range of those determined using multiple electrodes^20–22^ or optical mapping, although the latter required reduction of excitability by high extracellular K^+^ concentrations^15,23^. Mild reduction of gap junctional coupling decreased λ by 22% and reduced I_thr_ by more than the double (46%). Similarly, PVC vulnerability in cardiac infarction was increased by almost 50%. These results underline the importance of electrical coupling in the prevention of PVCs^24^ and are in line with increased arrhythmogenesis in cardiac fibrosis or after myocardial infarction by reducing the electrical sink of the surrounding myocardium^1,25^. In addition, the infarct led to RMP depolarization of ∼3.5 mV in the border zone, which by itself would increase PVC vulnerability by only ∼20% (**Fig. 3e**). Thus, depolarization explains <50% and loss of electrical sink >50% of the increased PVC vulnerability in the border zone.

Beyond diastolic values, we were able to determine R_m_ and λ during the cardiac cycle using triggered optogenetic current clamp. R_m_ was lowest in the diastole reflecting the importance of I_K1_ for preventing diastolic PVCs and highest during early repolarization indicating a vulnerability to PVCs in this phase. Quantitatively, R_m_ was ∼2-fold larger during early repolarization compared to diastole (**Fig. 1k**), which is less than the difference described in single cardiomyocytes (∼200-fold)^18,26^. This discrepancy may be due to non-synchronous repolarization in intact hearts, which can reduce R_m_ during repolarization because some cardiomyocytes are still in plateau or already at diastole, dynamically acting as electrical sink in a well-coupled syncytium, an effect further promoted by the increased λ at early repolarization (**Fig. 6e**). Furthermore, increased electrical coupling during repolarization reduces gradients of repolarization and, accordingly, lowers the chance that a PVC induces reentry during this highly vulnerable phase in which R_m_ is high.

Heart failure and myocardial infarction are associated with increased PVC susceptibility^5–7,27^ and ∼4-9 mV depolarization of RMP^28,29^, which has been attributed to reduced I_K1_. Partial block of I_K1_ by 10 µM BaCl_2_, which was shown to reduce I_K1_ similarly as in heart failure^27^, did not depolarize RMP (**Fig. 2e**) in our experiments, presumably because RMP was close to the K^+^ reversal potential. Thus, the RMP depolarization seen in heart failure may be rather due to pathological depolarizing diastolic leak currents, which could result from a leaky mode of the Na^+^-K^+^-ATPase at reduced ATP levels^30–33^, from a sustained, TTX-sensitive Na^+^ current recorded during hypoxia^32,34,35^, or from Connexin hemichannels activated by ischemia^36^. To quantify excitability in response to RMP depolarization, we experimentally added a leak current by diastolic, low intensity illumination. Interestingly, leak-induced depolarization increased R_m_, potentially due to voltage-dependent polyamine block of I_K1_^3^. Quantitatively, the effect of 7 mV depolarization (**Fig. 3c**) was comparable to the effect of pharmacological I_K1_ reduction (**Fig. 2d**) but with much higher I_thr_ reduction (**Fig. 2k, 3e**). This suggests that RMP depolarization increases PVC vulnerability not only by increasing R_m_ but also by reducing the distance from RMP to threshold for AP initiation.

Comparing experimentally measured relationships between RMP, R_m_, and I_thr_ with mathematical simulations showed that the increase in R_m_ due to a depolarizing leak current was largely overestimated by the original model (**Fig. 4e**). This discrepancy could only be resolved by reducing I_K1_ inward rectification, but not by up- or downscaling of current densities (**Fig. 4d**). Thus, optogenetic current clamp allows quantification of I_K1_ inward rectification in the intact heart, which before was only possible by patch clamp of single cells. Kir2.1 is the main isoform of I_K1_ in mouse ventricle^37^ and its inward rectification was much stronger in isolated mouse cardiomyocytes (ir ∼0.04-0.09)^12,38,39^ and heterologous expression systems (ir ∼0.09)^40^ than in our experiments in the intact mouse heart (ir = 0.033, **Fig. 4d**). This could be due to higher ATP levels in Langendorff-perfused hearts (8 mM)^41^ compared to patch clamp solutions (2-3 mM)^12,39^, which binds polyamines and thereby reduces I_K1_ rectification^42^. Our simulations show the importance of low I_K1_ rectification to reduce cardiac excitability (**Fig. 4c, 5g**) and we speculate that low ATP levels in cardiac disease may abolish this protective mechanism. Thus, additionally to increase I_K1_ density, suggested before to treat arrhythmia^43–45^, we propose reduction of its inward rectification e.g. by direct interaction with the channel or by reducing ornithine decarboxylase activity to lower polyamine levels as a novel therapeutic concept preventing PVCs. It was shown that the Na^+^ channel blocker Flecainide also reduces Kir2.1 rectification^46^ and we propose that only optogenetic current clamp can measure the integrated effects of such multi-channel drugs on cardiac excitability in the intact heart.

We conclude that characterizing pathological pro-arrhythmic conditions and anti-arrhythmic therapies by analyzing their effects on R_m_ and I_thr_ (**Fig. 5g**) provides a more comprehensive assessment compared to approaches focusing only on effects on ion channels. Our optogenetic current clamp method is feasible in both computational simulations^47–49^ and experimental preparations, although the latter is currently limited to transgenic mouse models. However, the development of larger animal models by AAV gene transfer^50^, combined with this new methodology, is important to improve our understanding of cardiac arrhythmias and testing new therapeutic strategies such as reduction of I_K1_ rectification.

## Methods

### Animal model

All experiments were in accordance with the European Guideline for animal experiments 2010/63/EU. Experiments for determination of RMP and afterdepolarization amplitude were performed with 11 male and 7 female CD-1 wildtype hearts. For all other experiments, 33 male and 43 female transgenic hearts from a previously established mouse line^9,47^ expressing ChR2 (H134R mutation) and eYFP under control of the chicken-β-actin promotor and backcrossed at least 10 generations on a CD-1 genetic background were used.

### Microelectrode recording

Mice were sacrificed by cervical dislocation. Hearts were explanted and retrogradely perfused in Langendorff configuration at a constant pressure of 110 cmH_2_O with Tyrode’s solution (in mM: 142 NaCl, 5.4 KCl, 1.8 CaCl_2_, 2 MgCl_2_, 10 glucose, and 10 HEPES, pH 7.4, adjusted with NaOH, 37°C, 100% O_2_) containing 10 µM Blebbistatin (Enzo Life Science, TargetMol) to inhibit contractions. The resulting flow was measured with a flow meter (SLF3S-1300F, Sensirion). Chemicals and reagents were purchased from Sigma Aldrich, if not specified otherwise. To measure membrane potential, sharp microelectrodes (15-40 MΩ, 1B100F-4, World Precision Instruments, pulled with P-1000 Micropipette Puller, Sutter Instrument) filled with 3M KCl were inserted into the left ventricular free wall and pushed forward through the myocardial wall using a fast piezoelectric actuator with position output (Sensapex uMp, ∼5-10 µm steps). The actual depth was determined considering the position output and penetration angle (45°) of the microelectrode. Membrane potential was recorded with a microelectrode amplifier (BA03S, NPI electronics) and a PowerLab recording system (PowerLab 8/30, LabChart software, ADInstruments, 10 kHz sampling rate). Cells with an AP amplitude smaller than 70 mV were excluded from analysis.

### Optogenetic stimulation

For optogenetic stimulation, patterned illumination of the left ventricular free wall was performed with a 465 nm LED (LEDMOD HP, Omicron Laserage) and a digital mirror device (Polygon400-G, Mightex) coupled to a macroscope (THT, Scimedia) with a 1x objective (MVPLAPO, Olympus, 0.25 n.a.). Light intensities were calibrated using a power meter (PM100A, S130C, Thorlabs). Both atria were removed to reduce spontaneous heart rate and allow for optogenetic pacing with a cycle length of 275 ms. The threshold for optogenetic pacing was measured by restricting the illumination to a circular area of 11.2 mm^2^ (**Fig. 2-5**) or 6.2 mm^2^ (**Fig. 6**) or a square area of ∼1 mm^2^ (**Fig. 7**) and applying an S1-S2 protocol consisting of six S1 light pulses at a cycle length of 275 ms (10 ms, ∼350 µW mm^-2^) followed by a S2 stimulus of variable delay and light intensity. The pacing threshold was defined by the lowest light intensity of the S2 pulse that allowed for successful pacing in five consecutive S1-S2 iterations. If not otherwise indicated, pacing thresholds were determined during diastole with a S2 interval of 265 ms. Optogenetic RMP depolarization (**Fig. 3**) was achieved by using low intensity light of a second 460 nm LED (LED Hub, Omicron Laserage) and a light guide collimated with a lens for uniform illumination of the left ventricular free wall.

### RMP determination

Wildtype hearts were electrically paced (1-10 mA, 0.5 ms, biphasic, cycle length 275 ms) via a monopolar Platinum/Iridium microelectrode (Science Products, PI2PT30.1B10) placed on the epicardium. If the sharp microelectrode touched the epicardial surface, the offset of the microelectrode amplifier was set to zero and the electrode resistance was measured. The electrode resistance was determined at each cell and recording was discarded if the electrode resistance changed >10 MΩ from the initial value, indicating a break or clotting of the tip. RMP was determined by using the peak analysis module of LabChart and averaging over 10 s.

### Patch clamp

Adult ventricular cardiomyocytes were enzymatically isolated from ChR2 transgenic mice as described previously^9,51^ and plated at low density on laminin-coated (0.1%) coverslips in external solution (in mM: 142 NaCl, 5.4 KCl, 1.8 CaCl_2_, 2 MgCl_2_, 10 glucose, and 10 HEPES, pH 7.4, adjusted with NaOH, 22°C). Patch-clamp recordings were made with an EPC10 amplifier and the Patchmaster software (Heka) using the whole cell configuration. Patch pipettes (3-5 MΩ) were filled with internal solution containing (in mM) 50 KCl, 80 K-Asparatate, 1 MgCl_2_, 3 MgATP, 10 EGTA, 10 HEPES, pH 7.2, adjusted with KOH. Light-induced ChR2 currents were measured in voltage clamp mode at -75.4 mV by illuminating with a lightguide-coupled 465 nm LED (LED Hub, Omicron Laserage) with attenuation filters through a 20x objective (S-Fluor, 0.75 n.a.) on Eclipse TI-2E microscope (Nikon) using high intensity illumination (315 µW mm^-2^, 5 ms) followed by a low intensity subthreshold light pulse (13.6 µW mm^-2^, 20 ms) with a delay of 200 ms.

### Protocol for induction of afterdepolarizations

To induce spontaneous afterdepolarizations, we used a stimulation protocol consisting of fast electrical pacing (cycle length of 70-140 ms for 40 cycles), followed by a break of 10 s where afterdepolarizations were analyzed. After the break, hearts were paced with a cycle length of 215 ms for at least 30 cycles before repeating the protocol. If the spontaneous heart rate was too fast to allow for analysis of afterdepolarizations, 10 µM Carbachol was applied (in 1 of 7 hearts). Moving average filtering over 100 ms was performed on all membrane potential traces prior to analysis of afterdepolarization amplitude.

### Model of cryoinfarction

To investigate cardiac excitability after acute myocardial infarction, an acute cryoinjury infarct model was used that generates precise infarction size and allows for a well-defined localization of the infarcted area. The cryoinfarction was produced by applying a liquid nitrogen cooled copper probe 3 times for 15 s to the apical part of the free left ventricular wall^52^. Pacing thresholds of 10 illumination pattern (∼1×1 mm) which were successively moved from base to apex (330-660 µm steps) were measured before and after cryoinfarction. PVC vulnerability was defined as the reciprocal of the pacing threshold and set to zero if the maximum light intensity (∼800 µW mm^-2^) did not lead to successful pacing.

### Simulation of channelrhodopsin-2 currents

ChR2 currents were calculated using a previously described ChR2 gating model^11^, which was implemented in MATLAB (2020b, Mathworks) and calibrated to match the peak current of the subthreshold light pulse measured in patch clamp experiments of adult cardiomyocytes expressing ChR2 (−0.45±0.03 pA pF^-1^ at 13.6 µW mm^-2^, n=7). The ChR2 current for calculation of R_m_ was converted in pA by assuming a membrane capacity of 100 pF per cardiomyocyte. To precisely calculate the ChR2 current during the cardiac cycle to determine I_thr_ and R_m_, the applied, calibrated light intensities and the averaged membrane potential trace of each experiment were used because the ChR2 photocurrents depend on the membrane potential (**Fig. 1e,h**). To compensate for offset changes during an experiment, membrane potential traces were shifted so that the diastolic potential matched the RMP measured under control conditions (−75.4±0.6 mV, n=30, N=3). If the RMP was optogenetically depolarized, the diastolic potential was additionally corrected by the constant light-induced RMP depolarization (**Fig. 3**). To account for peak current desensitization, the simulations started with 15 s of pre-pacing (membrane potential trace of AP without subthreshold illumination, cycle length 275 ms) before the AP cycle of interest was simulated.

### Computational simulation of cardiac electrophysiology

Single cell simulations were performed using a murine ventricular cardiomyocyte model^12^. Computational experiments were conducted using the openCARP simulation environment (version 8.2)^53^. To allow for optogenetic determination of R_m_ and pacing threshold, the calibrated ChR2 gating model described above^11^ was integrated into the cardiomyocyte model. Analogous to the experiments, R_m_ was determined in the simulations via delivery of a subthreshold light pulse (20 ms, 15 µW mm^-2^) after simulating at least 10 AP cycles (cycle length 275 ms) induced by optogenetic pacing. The pacing threshold was defined as lowest light intensity at which 10 of 10 light pulses (10 ms) induced APs. RMP was defined as the minimum membrane potential after simulating 10 AP cycles. The RMP was depolarized by adding a cation background current (I_leak_) to the cardiomyocyte model, mimicking the ChR2 current in the experiments with a K^+^ conductance of 0.5 relative to Na^+^ conductance^54^. I_leak_ was calculated as:

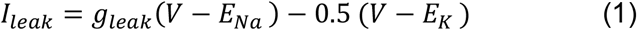

with V the membrane potential, g_leak_ a scaling factor, and E_Na_/E_K_ the Na^+^/K^+^ reversal potentials, respectively. Simulations for **Fig. 4a-d** were performed without modification of the I_K1_ inward rectification parameter (ir). The experimentally measured increase of R_m_ (8.7% at 3.5 mV RMP depolarization) could be achieved by reducing ir to 37% (from initial 0.0896 to 0.0332) in the I_K1_ equation of the model, which was subsequently used (**Fig. 4e-g, Fig. 5d-g**):

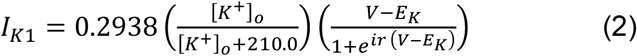

Values of R_m_, RMP, and I_thr_ were only used if the model was stable (i.e., the simulated cell returned to RMP after each AP). Calculation of percentage changes of pacing thresholds were based on ChR2 photocurrents in pA pF^-1^. Simulation of the effect of the multi-channel blockers Amiodarone and Dronedarone were adapted from previously described half maximal inhibitory concentrations and Hill coefficients^13^.

The openCARP code for determining R_m_, I_thr_ and R_m_ change in response to 3.5 mV RMP depolarization induced by I_leak_ is available as supplementary material.

### Data analysis and statistics

The analysis of the data, including filtering, calculation of ChR2 current and R_m_, detection of different AP phases and parameters, determination of afterdepolarization amplitude, and fitting of λ by the method of least squares was performed with MATLAB R2020b with the optimization toolbox (MathWorks). For analysis of cycle-dependent R_m_, the diastole was defined between 200-240 ms after AP initiation, early and late repolarization as timepoints of 70% (APD70) and 90% (APD90) of repolarization, respectively, and plateau timing as 0.5*APD70. To determine R_m_ and pacing threshold at 3.5 and 7 mV RMP depolarization, data were linearly interpolated (**Fig. 3c,e**). λ was obtained by fitting the measured subthreshold membrane potential change of different light pattern with

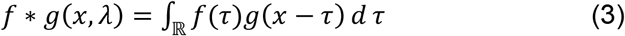

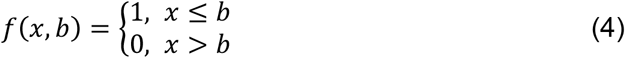

is the function of light depending on the position of the microelectrode *b* and

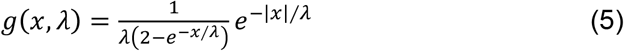

the weight function. Fits of λ with a mean squared error larger than 0.3 mm^2^ were excluded from analysis. Cycle-dependent λ was measured 200 ms after AP initiation (diastole), at APD70

## Supporting information

OpenCARP code used in the manuscript

## Ethical Approval

not applicable

## Authors’ contributions

JSL and PS conceptualized the project and wrote the manuscript. JSL performed and analyzed experiments using intact hearts. JSL and PMB designed the computational framework and performed simulations using openCARP. DM performed patch clamp experiments.

## Availability of data and materials

Original data will be provided upon personal and reasonable request to the corresponding author.

## Acknowledgments

We thank F. Holst (University of Bonn) for technical assistance, M. Vaisband and J. Hasenauer (University of Bonn) for fitting of the relationship between R_m_ and λ.

## Abbreviations

PVC: premature ventricular contraction
I_K1_: inwardly rectifying K^+^ current
RMP: resting membrane potential
R_m_: input resistance
ChR2: channelrhodopsin-2
I_ChR2_: channelrhodopsin-2 current
λ: length constant
AP: action potential
APDx: action potential duration at percentage of repolarization
ΔE: change in membrane potential
DAD: delayed afterdepolarization
I_NaK_: current of the Na^+^-K^+^-ATPase
I_Nab_: sodium background current
I_Na_: voltage-dependent sodium current
Ir: inward rectification of I_K1_
I_leak_: background leak current
MI: myocardial infarction

