## Supplementary material for "Optogenetic quantification of source sink relationship in intact hearts to explain cardiac arrhythmia initiation and protection": OpenCARP code used in the manuscript: Calculate_PT_RMP_RM_WithGLeakAdjust.sh

```
#!/bin/bash -u

DEBUG=0;

# WARNING: Both of these variables must be set by the user based on machine-specific settings

BENCH=/Software/openCARP/v8.2/bin/bench.opt;
IMPSO=/data/Projects/Langen_OptogeneticCurrentClamp/repo/MurineMouse_leak_ik1.so;

# WARNING: Once machine-specific settings have been applied above, comment out the next two lines:
echo "error: environment variables need to be set; please edit the script" >&2; 
exit 1;

# for Rm determination subthreshold pulse of 15 µW/mm2

GNaB=$1;       shift; # default value: 1.0
GNaK=$1;       shift; # default value: 1.0
GK1=$1;        shift; # default value: 1.0
SPERMIN=$1;    shift; # default value: 1.0
LeakFactor=$1; shift; # default value: 0.75
DepolTarget=$1;shift; # default value: 3.5

# IChR2 parameters, as in Williams, Xu, et al.
GCHR2=0.4;
GAM=0.1; 
TEMP=37;

BENCH="$BENCH --dt=0.002 --load-module=$IMPSO --imp=MurineMouse_leak_ik1 --plug-in=IChR2 --plug-par=gam=$GAM,gChR2=$GCHR2,Temp=$TEMP";
OUTDIR=Calculate1_PT_RMP_RM15_GNaBx${GNaB}_GNaKx${GNaK}_GK1x${GK1}_Spermin${SPERMIN}_LeakFactor${LeakFactor}_DepolTarget${DepolTarget};
rm -f ${OUTDIR}.log;
mkdir -p $OUTDIR;

GLeak=0.0;
BENCHCMD="$BENCH --imp-par GNaKLeak*${GLeak},i_NaKLeak_K_factor*${LeakFactor},g_K1*${GK1},g_Nab*${GNaB},i_NaK_max*${GNaK},spermin*${SPERMIN}";

EQDUR=1000;
FOUT=$OUTDIR/Part1_Equilibrate${EQDUR};
echo "Part 1: Identify initial RMP (dur = $EQDUR ms)" >> $OUTDIR.log;
if [[ $DEBUG == "1" ]] || [[ ! -e ${FOUT}.txt ]] || [[ ! -e ${FOUT}.sv ]]; then
    $BENCHCMD \
        --stim-curr=0 \
        --light-irrad=0 \
	--duration=$EQDUR \
        --save-ini-time=$EQDUR \
        --save-ini-file=${FOUT}.sv \
	--fout=${FOUT} \
        &> ${FOUT}.log;
    mv Trace_0.dat ${OUTDIR}/Part1_Equilibrate${EQDUR}_Trace.dat;
fi;

RMP_INIT=$(tail -n1 $FOUT.txt |awk '{print(1*$2)}');
RMP_TARGET=$(echo "scale=40; $RMP_INIT + $DepolTarget" |bc);

echo "Part 2: Calibrate to target RMP ($RMP_INIT + $DepolTarget = $RMP_TARGET)" >> $OUTDIR.log;

function bisect { echo "scale=40; $1+0.5*($2-$1)" |bc; };

RMP_TOL=0.01;
GLeak_LO=0.0;
GLeak_HI=0.1;
GLeak_HI_OK=0;
N=0;
mkdir -p $OUTDIR/Part2_AdjustGLeak${RMP_TOL};
while [[ 1 ]]; do
    NN=$(printf %03d $N);
    if [[ $GLeak_HI_OK == "0" ]]; then
        GLeak=$GLeak_HI;
    else
        GLeak=$(bisect $GLeak_LO $GLeak_HI);
    fi;
    BENCHCMD="$BENCH --imp-par GNaKLeak*${GLeak},i_NaKLeak_K_factor*${LeakFactor},g_K1*${GK1},g_Nab*${GNaB},i_NaK_max*${GNaK},spermin*${SPERMIN}";

    FOUT=$OUTDIR/Part2_AdjustGLeak${RMP_TOL}/${NN}_Test${GLeak};
    if [[ $DEBUG == "1" ]] || [[ ! -e ${FOUT}.txt ]] || [[ ! -e ${FOUT}.sv ]]; then
        $BENCHCMD \
            --stim-curr=0 \
            --light-irrad=0 \
	    --duration=$EQDUR \
            --save-ini-time=$EQDUR \
            --save-ini-file=${FOUT}.sv \
	    --fout=${FOUT} \
            &> ${FOUT}.log;
        mv Trace_0.dat $OUTDIR/Part2_AdjustGLeak${RMP_TOL}/${NN}_Test${GLeak}_Trace.dat;
    fi;
    RMP=$(tail -n1 $FOUT.txt |awk '{print(1*$2)}');
    echo "Part 2: Calibrate to target RMP (test[$N] GLeak = $GLeak nS/pF): RMP = $RMP mV)" >> $OUTDIR.log;

    if [[ $(echo $RMP $RMP_TARGET |awk '{if($1<$2) print(1); else print(0);}') == "1" ]]; then
        if [[ $GLeak_HI_OK == "0" ]]; then
            GLeak_HI=$(echo "scale=40; $GLeak_HI*2" |bc);
        else
            GLeak_LO=$GLeak;
        fi;
    else
        GLeak_HI_OK=1;
        GLeak_HI=$GLeak;
    fi;

    N=$((N+1));

    if [[ $(echo $RMP $RMP_TARGET $RMP_TOL |awk '{diff=$2-$1; if(diff<0) diff=-diff; if(diff<$3)print(1); else print(0);}') == "1" ]]; then break; fi;
done;

EQUILIBRIUM_SV_FOR_DESIRED_RMP=${FOUT}.sv;

EETOL=0.0001;
EELO=0.0;
EEHI=0.01; # 0.8
EEHI_OK=0;
N=0;
mkdir -p $OUTDIR/Part3_FindPT${EETOL};
while [[ 1 ]]; do
    NN=$(printf %03d $N);
    EE=$(bisect $EELO $EEHI);
    
    if [[ $(echo $EE | awk '{if($1>2.0) print(1); else print(0);}') == "1" ]]; then
        echo "$OUTDIR [Aborted] Could not detect pacing threshold. Repolarization failure?" |tee $OUTDIR/OutputParams.txt;
        exit;
    fi;
    
    FOUT=$OUTDIR/Part3_FindPT${EETOL}/${NN}_Test${EE};
    if [[ $DEBUG == "1" ]] || [[ ! -e ${FOUT}.txt ]] || [[ ! -e ${FOUT}.sv ]]; then
        $BENCHCMD \
            --read-ini-file=$EQUILIBRIUM_SV_FOR_DESIRED_RMP \
            --stim-curr=0 \
            --light-irrad=$EE \
            --light-dur=10 \
            --light-numstim=10 \
            --light-bcl=275 \
            --light-start=50 \
	    --duration=3000 \
            --save-ini-time=3000 \
            --save-ini-file=${FOUT}.sv \
            --fout=${FOUT} \
            &> ${FOUT}.log;
        mv Trace_0.dat $OUTDIR/Part3_FindPT${EETOL}/${NN}_Test${EE}_Trace.dat;
    fi;
    NZC=$(cat $FOUT.txt |awk 'BEGIN{zc=0;}{v=$2; if(NR>1&&vp<0&&v>=0)zc++; vp=v;}END{print(zc);}');
    echo "Part 3: Pacing Threshold (test[$N] Ee = $EE mW/mm^2): # zero crossings = $NZC)" >> $OUTDIR.log;

    if [[ $NZC -lt 10 ]]; then
        if [[ $EEHI_OK == "0" ]]; then
            EEHI=$(echo "scale=40; $EEHI*2" |bc);
        else
            EELO=$EE;
        fi;
    else
        EEHI_OK=1;
        EEHI=$EE;
        FOUT_OK=$FOUT;
    fi;

    N=$((N+1));

    if [[ $(echo $EELO $EEHI $EETOL|awk '{if(($2-$1)<$3)print(1); else print(0);}') == "1" ]]; then break; fi;
done;

PT=$EEHI;
RMP=$(tail -n1 $FOUT_OK.txt |awk '{print(1*$2)}');
echo "Part 3: Pacing Threshold = $PT mW/mm^2 (RMP = $RMP mV)" >> $OUTDIR.log;

EE=$(echo "scale=40; 1.2*${PT}" |bc);
FOUT=$OUTDIR/Part3_FindPT${EETOL}_1.2xThresh;
echo "Part 3(a): Final run with 1.2x Thresh (Ee = $EE mW/mm^2)" >> $OUTDIR.log;
if [[ $DEBUG == "1" ]] || [[ ! -e ${FOUT}.txt ]] || [[ ! -e ${FOUT}.sv ]]; then
    $BENCHCMD \
        --read-ini-file=$OUTDIR/Part1_Equilibrate${EQDUR}.sv \
        --stim-curr=0 \
        --light-irrad=$EE \
        --light-dur=10 \
        --light-numstim=10 \
        --light-bcl=275 \
        --light-start=50 \
	--duration=3000 \
        --save-ini-time=2725 \
        --save-ini-file=${FOUT}.sv \
        --fout=${FOUT} \
        &> ${FOUT}.log;
    mv Trace_0.dat $OUTDIR/Part3_FindPT${EETOL}_1.2xThresh_Trace.dat;
fi;

EE=$(echo "scale=40; 0.1*${PT}" |bc);
FOUT=$OUTDIR/Part4_TestPulse_+200ms;
if [[ $DEBUG == "1" ]] || [[ ! -e ${FOUT}.txt ]] || [[ ! -e ${FOUT}.sv ]]; then
    $BENCHCMD \
        --read-ini-file=$OUTDIR/Part3_FindPT${EETOL}_1.2xThresh.sv \
        --stim-curr=0 \
        --light-irrad=0.015 \
        --light-dur=20 \
        --light-numstim=1 \
        --light-start=0 \
	--duration=100 \
        --save-ini-time=100 \
        --save-ini-file=${FOUT}.sv \
        --fout=${FOUT} \
        &> ${FOUT}.log;
    mv Trace_0.dat $OUTDIR/Part4_TestPulse_+200ms_Trace.dat;
fi;

# Workaround to figure out which column of Trace_0.dat contains the IChR2 values:
COLIDX=$(cat MurineMouse_leak_ik1_trace_header.txt IChR2_trace_header.txt |awk '{if($1=="IChR2")print(1+NR);}');

IDIFF=$(cat $OUTDIR/Part4_TestPulse_+200ms_Trace.dat |awk 'BEGIN{tt=20;}{t=$1;IChR2=$'$COLIDX'; dt=t-tt; if(dt<0)dt=-dt; if(dt<0.1) {print(1*IChR2);exit;}}');
VDIFF=$(cat $FOUT.txt |awk '{if(NR==1){v0=v1=$2;} if($2>v1) v1=$2;}END{print(v1-v0)}');
RM=$(echo $VDIFF $IDIFF |awk '{print(-10*$1/$2);}');
echo "Part 4: Run with 0.1x Thresh (Ee = $EE mW/mm^2) -- R_m = $RM MΩ (ΔV_m = $VDIFF mV)" >> $OUTDIR.log;

echo "$OUTDIR $PT $RMP $RM $VDIFF" |tee $OUTDIR/OutputParams.txt;

exit 0;
```
