## Supplementary material for "Optogenetic quantification of source sink relationship in intact hearts to explain cardiac arrhythmia initiation and protection": OpenCARP code used in the manuscript: Reproduce_FigS1.sh

```
#!/bin/bash -u

# Script to reproduce Fig. S1 from the following paper:
# Loewe A, Lutz Y, Wilhelms M, Sinnecker D, Barthel P,
#   Scholz EP, Dössel O, Schmidt G, Seemann G. In-silico
#   assessment of the dynamic effects of amiodarone and
#   dronedarone on human atrial patho-electrophysiology.
#   Europace. 2014 Nov;16 Suppl 4:iv30-iv38.
#   doi: 10.1093/europace/euu230. PMID: 25362168.

# Final results can be visualized by running:
# $ xmgrace Amiodarone.agr
# $ xmgrace Dronedarone.agr

VAR[1]="IKr";
VAR[2]="IKs";
VAR[3]="INa";
VAR[4]="ICaL";
VAR[5]="INaCa";
VAR[6]="INaK";

IC50[1]=2.80;
IC50[2]=3.84;
IC50[3]=4.84;
IC50[4]=5.80;
IC50[5]=3.30;
IC50[6]=15.60;

nH[1]=0.91;
nH[2]=0.63;
nH[3]=0.76;
nH[4]=1.00;
nH[5]=1.00;
nH[6]=1.00;

for N in {1..6}; do 
    for D_AMIO in \
        0.2300 0.3196 0.4441 0.6170 0.8573  1.1913  1.6553  2.3000 \
               3.1958 4.4406 6.1702 8.5735 11.9128 16.5528 23.0000; do
        echo $D_AMIO |awk '{print($1, (1+('${IC50[$N]}'/$1)^'${nH[$N]}')^-1)}';
    done > Amiodarone_scf_${N}_${VAR[$N]}.dat;
done;

VAR[1]="IKr";
VAR[2]="IKur";
VAR[3]="IKs";
VAR[4]="INa";
VAR[5]="ICaL";

IC50[1]=0.0591;
IC50[2]=1.00;
IC50[3]=5.60;
IC50[4]=0.54;
IC50[5]=0.83;

nH[1]=0.80;
nH[2]=1.00;
nH[3]=0.51;
nH[4]=2.03;
nH[5]=2.75;

for N in {1..5}; do 
    for D_DRON in \
        0.0210 0.0292 0.0405 0.0563 0.0783 0.1088 0.1511 0.2100 \
               0.2918 0.4054 0.5634 0.7828 1.0877 1.5113 2.1000; do
        echo $D_DRON |awk '{print($1, (1+('${IC50[$N]}'/$1)^'${nH[$N]}')^-1)}';
    done > Dronedarone_scf_${N}_${VAR[$N]}.dat;
done;

exit 0;
```
