## Supplementary material for "Optogenetic quantification of source sink relationship in intact hearts to explain cardiac arrhythmia initiation and protection": OpenCARP code used in the manuscript: README.md

This README file contains basic information to use openCARP parameter files
and driver scripts that accompany the publication by Langen, Boyle, Malan,
and Sasse, first published as a biorXiv preprint entitled "Optogenetic 
quantification of source sink relationship in intact hearts to explain 
cardiac arrhythmia initiation and protection" in August, 2024.

If you reuse this code, please cite the most recent version of this
publication (either the preprint or the final published version, as
appropriate).

For help, please consider emailing Dr. Patrick Boyle at the University of
Washington.

##############################################################################
Important prerequisites and setup steps:
##############################################################################

1) To run this file, you must have a working version of openCARP compiled from
source code. At the time of compilation, the flag to "enable ionic shared 
library loading" must be asserted. For example, this my_switches.def file:

#DEBUG=1                # compile a debug version
#PROFILE=1              # turn on standard gcc profiling
#SCALASCA_INSTRUMENT=1  # instrument code with Scalasca profiling
#SILENT_MAKE=1          # embellish make output
DLOPEN=1 # enable ionic shared library loading
MAKE_FLG="-j" # activate parallel make
#TAGEXE=etags -a        # generate tags
#USE_SIMD = avx2        # version of support for Intel vectorizing SIMD code
#USE_OPENMP=1            # use OpenMP
#CVODE=1                # use CVODE 

Note that DLOPEN=1 has been uncommented.

These scripts have been extensively tested with v8.2 of the openCARP code,
which should be available via the git repository (i.e., immediately after
cloning a clean download of the source, run git checkout v8.2, then compile).

2) Once you have a working version of openCARP, navigate to the directory
where the MurineMouse_leak_ik1.model file packaged with this archive is
located (e.g., /data/Projects/Langen_OptogeneticCurrentClamp), then use the
openCARP .model compilation tool to create a corresponding .so file:

$ make_dynamic_model.sh MurineMouse_leak_ik1.model
/Software/openCARP/v8.2/physics/limpet/src/python/limpet_fe.py MurineMouse_leak_ik1.model /Software/openCARP/v8.2/physics/limpet/models/imp_list.txt .
/Software/openCARP/v8.2/physics/limpet/src/python/make_dynamic_model.py --compiler=mpicc --cflags=-fPIC -shared -std=c++11 -lm -I/Software/openCARP/v8.2/physics/limpet/src -I/Software/openCARP/v8.2/numerics -I/Software/openCARP/v8.2/simulator -I/Software/openCARP/v8.2/fem -I/Software/openCARP/v8.2/fem/slimfem/src -I/Software/openCARP/v8.2/param/include -I/usr/lib/petsc//include -I/usr/lib/petsc///include --dynamic_prefix= MurineMouse_leak_ik1.model
Dynamic model generator
Generating: MurineMouse_leak_ik1 -> /tmp/MurineMouse_leak_ik1_dyn.cc
mpicc -fPIC -shared -std=c++11 -lm -I/Software/openCARP/v8.2/physics/limpet/src -I/Software/openCARP/v8.2/numerics -I/Software/openCARP/v8.2/simulator -I/Software/openCARP/v8.2/fem -I/Software/openCARP/v8.2/fem/slimfem/src -I/Software/openCARP/v8.2/param/include -I/usr/lib/petsc//include -I/usr/lib/petsc///include -I/tmp  -o /tmp/MurineMouse_leak_ik1.so /tmp/MurineMouse_leak_ik1.cc /tmp/MurineMouse_leak_ik1_dyn.cc

3) Before running any of the driver scripts, be sure to edit the following
three files:
- Experiment01_AdjustGLeak/Calculate_PT_RMP_RM_WithGLeakAdjust.sh
- Experiment02_CharacterizeExcitability/Calculate_PT_RMP_RM.sh
- Experiment03_AmiodaroneDronedarone/Calculate_PT_RMP_RM_WithDrugs.sh

All three files contain the following two environment variable assignments:
BENCH=/Software/openCARP/v8.2/bin/bench.opt;
IMPSO=/data/Projects/Langen_OptogeneticCurrentClamp/MurineMouse_leak_ik1.so;

These must be updated to match the absolute paths to the openCARP source
directory and the working directory for these simulations on your machine. If
these changes are not made, the driver scripts will fail with an eror.

##############################################################################
Basic instructions for usage
##############################################################################

Once the prerequisite and setup steps are complete, navigate to one of the
three directories:
- Langen_OptogeneticCurrentClamp/repo/Experiment01_AdjustGLeak
- Langen_OptogeneticCurrentClamp/repo/Experiment02_CharacterizeExcitability
- Langen_OptogeneticCurrentClamp/repo/Experiment03_AmiodaroneDronedarone

Then, run the test harness script as follows:
- bash -u Exp[XX]_Run_TestHarness.sh Exp[XX]_TestHarness

(Modify the [XX] section as needed for the subdirectory as needed)

If problems are encountered, see the next section for helpful tips.

##############################################################################
Tips for debugging
##############################################################################

- All three of the Calculate_[...].sh scripts have debug modes that can be
enabled by changing the header from DEBUG=0 to DEBUG=1. This will ensure that
simulations are run no matter what, even if their output files already exist.

- Before running a particular test harness, it is a good idea to run a single
iteration of the corresponding Calculate_[...].sh script with basic settings,
which can be obtained by examining the first line of Exp[XX]_TestHarness.dat
in each directory. For example:

$ bash -u Calculate_PT_RMP_RM.sh 1.0 1.0 1.0 1.0 0.0 1.5

- To temporarily redirect script output for the Calculate_[...].sh scripts to
the command line (instead of an output log), find-replace:
`` >> $OUTDIR.log ''
with
`` |tee -a $OUTDIR.log ''

- In all three Calculate_[...].sh scripts, there is a line where the ChR2
current differential is calculated (IDIFF="[...]"). This one-liner uses a
shortcut that exploits the way openCARP outputs data for simulations with an
ionic model (in this case, the Murine Mouse model) is augmented with a plug-
in (in this case, the Williams et al. model of ChR2). The code as published
uses a trick to find this column; other versions of openCARP may misbehave.

If your execution of the script produces nonsense values, it may be because
the way openCARP handles state variables has changed in your version of the
code. This should not be an issue for users who follow the instructions above
(i.e., use openCARP v8.2). Nevertheless, if this appears to be an issue,
please run the following command to discover which column of the Trace_0.dat
files have IChR2 information:

$ cat MurineMouse_leak_ik1_trace_header.txt IChR2_trace_header.txt |awk '{print(1+NR,$1)}' |column -t
2   E_K
3   V
[.....]         <== extra output removed for brevity's sake
22  sv->C2
23  IChR2       <== in this case, n = 23 is correct
24  IChR2_tot
25  O1
[.....]
